# Stimulation of the Frontal Aslant Tract’s origin in the caudal superior frontal gyrus alters ongoing spontaneous rhythmic activity independently from the effector. Evidence from tractography-guided Transcranial Magnetic Stimulation

**DOI:** 10.1101/2024.11.21.624661

**Authors:** Marco Tagliaferri, Lea Glaubig, Valentina DiChiaro, Luigi Cattaneo

**Author notes:** **Corresponding author:** Luigi Cattaneo., University of Trento, Center for Mind/Brain Sciences., Via delle Regole 101.

## Abstract

the crown of the human Superior Frontal Gyrus (SFG-crown) is a functionally independent region, nestled between the dorsal premotor and supplementary motor cortices, that supports internally-timed action control. The unique SFG-crown’s connectivity fingerprint by the Frontal Aslant Tract (FAT), suggests a caudal-rostral pattern of increasing abstractness of action representations. Coherently, since the mid-portion of the caudal SFG contains a representation of action strategies that involve internal timing, we hypothesized that the caudal portion of the SFG may be involved in action execution of internally-timed actions. To test this, we asked 21 healthy participants to perform a self-paced tapping movement with the right index finger or a self-paced articulation of the syllable /da/ while applying online single-pulse TMS to the posterior, middle and anterior origins of the FAT in the left SFG-crown. Results showed that effective TMS (compared to sham) impacted rhythm production in both tasks, only when applied to the posterior SFG region, by reducing the probability of motor events in the 200 ms following TMS. The present data support the hypothesis, that the posterior SFG-crown, associated with the most posterior origin of the FAT fibers, is involved in the production of internally-timed actions, in an effector-independent modality, suggesting a domain-general role in the execution of internally-timed movements.

## 1. INTRODUCTION

### 1.1. The caudal part of the convexity of the superior frontal gyrus (SFG)

is a cortical region of unclear functional attribution in humans. If considered along a caudal-cranial dimension, it contains at least two distinct cytoarchitectonic regions: Brodmann’s Area 6 and dorsal Brodmann’s area 8 or area 8A (Petrides and Pandya 1999; Glasser et al. 2016) and therefore contains the transition between the dysgranular premotor cortex (BA6) and the granular prefrontal cortex (BA8) (Geyer 2004). However, a parcellation with even finer granularity found in the Julich cortical atlas (Amunts et al. 2020) identifies 3 distinct regions along the SFG convexity: 6d1, 6d2 and 8d1. Similarly, Glasser’s atlas (Glasser et al. 2016) proposes 4 parcels within the BA6-BA8 complex: 6d, 6a, i6-8 and 8Ad. Along the medial-lateral dimension, recent evidence points at a subdivision of the caudal SFG in 3, rather than 2, distinct regions. Namely, in a medial-lateral order: a) the medial surface, b) the crown (or convexity) and c) the medial bank of the superior frontal sulcus (SFS). The medial surface by the presence of the supplementary / pre-supplementary motor areas (SMA and pre-SMA, hereafter referred to as SMA complex) and the medial BA8 (Picard and Strick 1996). On the lateral surface, the convexity of the SFG and the medial bank of the superior frontal sulcus (SFS) represent two separate identities. Indeed, the Julich atlas identifies several regions along the medial wall of the SFG that are distinct from those on the convexity: Areas 6d3 and 8d2. Similarly, the Glasser atlas draws a separation between the convexity of SFG and the medial bank of the SFS. Summing up, the caudal SFG seemingly contains 3 separate regions: the medial wall, the crown and the medial bank of the SFS. The medial wall is very well characterized functionally as SMA-complex. Similarly, the medial bank of the SFG is what is commonly referred to as dorsal premotor cortex, because the core region of the PMd that contains upper limb movements seems to be localized inside the SFS (Germann et al. 2005; Amiez et al. 2006; Chouinard and Paus 2006; Mayka et al. 2006; Tomassini et al. 2007) rather than on the convexity of the SFG. Nestled between the PMd and the SMA-complex, lies the convexity of the SFG, which is poorly defined from a functional point of view.

### 1.2. The connectional fingerprint of the SFG

is potentially of great value in cracking the functional parcellation of otherwise anatomically uniform region. The Frontal Aslant Tract is a white matter bundle that connects the SFG-crown with the inferior frontal gyrus (IFG). Unlike previously thought, the dorsal origin of the FAT is not in the medial wall, but rather in the SFG convexity (Tagliaferri et al. 2024). Interestingly, the FAT fibers run mostly parallel, with very little divergence /convergence between the dorsal (SFG) and the ventral (IFG) origins. As a consequence there is a strict correspondence between the SFG and the IFG sub-regions that are connected by FAT sub-bundles. The IFG origins of the FAT correspond, in a caudal-cranial order, to 3 well-defined sub-regions: ventral Brodmann’s area 6 (BA6), BA44 and BA45. (Briggs et al. 2020; Tagliaferri et al. 2024) These have a well-established functional hierarchy, becoming more complex in more cranial regions, representing orderly: movements, then actions and then the relation between actions and their meaning. For example, for what concerns linguistic processes in the left IFG, BA6 contains phonological representations, BA44 lexical representations and BA45 semantic and grammatical representations (see for example (Gough et al. 2005; Ishkhanyan et al. 2020)). The FAT connectivity allows us to hypothesize a similar hierarchy in the corresponding FAT origin regions in the SFG, with movements in the posterior part, actions in the middle part and meaning of action/relations between actions in the rostral portion. The individual functional specialization of the IFG and of the SFG can be hypothesized, according to the non-human and human primate literature, to be that of internally-generated behavior in the SFG versus externally-triggered behavior in the IFG. Indeed, neurophysiological findings in the monkey describe the ventral premotor regions representing 1:1 sensorimotor, bottom-up associations, and the dorsal and medial premotor regions specialized in internally-generated and conditional actions (Hoshi and Tanji 2004, 2006; Mita et al. 2009) and common models of human frontal lobe function show a medial-lateral gradient with internally controlled behavior represented medially and bottom-up sensorimotor behavior represented laterally (O’Reilly 2010; Stuss 2011).

### 1.3. The hypothesis of IFG-SFG homology

was recently validated in part, by showing that homologous regions in the IFG and SFG, connected by FAT fibers perform similar computations and are in direct competition for overt behavior. (Tagliaferri et al. 2023) We observed that in a pre-cued reaction time task, participants could perform the button press either by using a predictive (moving on the basis of an anticipatory, internal time estimation of the onset of the GO-cue) or a reactive (waiting for the GO-cue to happen and then moving) strategies. The two strategies are mutually incompatible and are chosen on a trial-by-trial basis. By using tractography-guided transcranial magnetic stimulation (TMS), we observed that stimulation of the origins of the middle part of the FAT, i.e. BA44 or *pars opercularis* of the IFG and the mid-SFG biased the choice towards a reactive or towards a predictive strategy, respectively. This finding was taken as evidence that the SFG and the IFG contain two opposite action strategies. The SFG represents actions based on internal timing, while the IFG represents actions based on external cues.

### 1.4. Current study hypothesis and aim

Summing up, the SFG-crown seems to represent internally-timed actions, and the middle portion of the SFG seems to represent entire action strategies. According to our hierarchical model, we therefore hypothesized that the posterior portion of the SFG-crown should be involved in the direct production of internally-timed movements. To test this hypothesis we performed an experiment using online TMS applied to the posterior, middle and anterior portions of the SFG, as defined by individual FAT connectivity. We chose simple internally-timed behavior, i.e. repetitive movements performed with either the upper limb (finger tapping) or with the orofacial effectors (repeated utterance of a syllable). We expected only TMS applied to the posterior SFG to interfere with the execution of the tasks. Regarding the two effectors used, we hypothesized two possible outcomes, i.e. that TMS effects could be effector-specific or generalized across effectors, thus indicating a domain-general role of the posterior SFG in setting the timing for internally-generated actions.

## 2. METHODS

### 2.1. Participants

Twenty-one participants (11 females and 10 males; age range 20–36 years) were recruited for this study. The experimental protocol was approved by the University of Trento’s local Ethical Committee (protocol number 2020_035), and informed consent was obtained from all subjects prior to participation. Before the experiment, all participants were screened for contraindications to transcranial magnetic stimulation (TMS) following established guidelines (Wassermann 1998; Rossi et al. 2009, 2021). Each participant attended two separate sessions: the first involved MRI-DWI scanning, and the second involved neuronavigated single-pulse TMS stimulations during the performance of two tasks. All of them experienced TMS stimulations over the left hemisphere, and ten of them a second identical session stimulating the right hemisphere.

### 2.2. Behavioral task

We asked 20 healthy human adult volunteers to perform in 2 separate sessions, two motor tasks involving A) self-paced repetitive speech (syllable) production or B) self-paced finger tapping. Sixty trials per condition (2 tasks x 4 stimulation conditions) were recorded for a total of 480 trials per participant and per each of the two tasks. The number of motor events (syllables or taps) was different between each single trial, since the timing of the events was completely left to the participant’s own initiative. Subjects were sitting in front of a desk with their head constrained in the lateral dimension by the arms of a optometrist’s-like chinrest, but without the chinrest part, because immobilizing the chin would have prevented the subjects from articulatory utterances. Stability of the scalp target of TMS was granted by constant monitoring of the neuronavigator system, which accounts for tilt and coil-scalp distance (cfr. further paragraphs).

### 2.3. T1w data preprocessing

Anatomical imaging was conducted using a 3T MAGNETOM Prisma scanner (Siemens Healthcare) equipped with a 64-channel head-neck RF receive coil. We acquired 3D T1-weighted images (multiecho-MPRAGE, 1 mm isotropic) and diffusion-weighted images (DWI; 2 mm isotropic, TE/TR = 76/4200 ms, with b-values of 0, 700, 1000, and 2850 s/mm^2^ across 32, 64, and 64 directions, respectively). Standard preprocessing was applied to the T1-weighted images for all participants. Initially, raw DICOM images were converted to NIfTI format using the dcm2niix package. The images were then aligned to the MNI152 T1-weighted template through rigid registration to ensure anterior-posterior commissure (AC-PC) alignment (Avants et al. 2008). Brain masks and segmentations were generated using MRtrix3 and the BET2 package (Tournier et al., MRtrix3), followed by N4 bias field correction. Additionally, the T1-weighted images were segmented into five brain tissue types using MRtrix3, and a synthetic T2-weighted image was produced using the AFNI toolkit in order to use it with TORTOISE toolkit during the diffusion imaging preprocessing.

### 2.4. DWI data preprocessing

The diffusion-weighted imaging data were preprocessed using the TORTOISE toolkit (Pierpaoli et al., 2010). Utilizing the synthetic T2-weighted image derived from the T1-weighted scan, we corrected for Gibbs ringing, thermal noise, eddy currents, and motion artifacts using the DIFFPREP tool. Susceptibility-induced echo-planar imaging (EPI) distortion was addressed with the DRBUDDI tool through diffeomorphic registration (Avants et al. 2008). This process incorporated both anterior-posterior and posterior-anterior phase-encoding directions of the DWI data, as well as information from an undistorted structural MRI. Bias field inhomogeneities were corrected using methods described by Tustison et al. (2010). Streamline tractography was performed using MRtrix3 software (Tournier et al., 2012). We applied multishell multitissue constrained spherical deconvolution (CSD; lmax = 6, threshold = 0.5) to estimate the white matter fiber orientation distribution functions (fODFs), using the Dhollander method for response function estimation (Dhollander et al. 2016). Deterministic tractography based on CSD was then computed with parameters set to a cutoff of 0.001, a maximum angle of 75°, a step size of 5 mm, and streamline lengths between 20 mm and 250 mm. Tractography was initiated with random seeding of 107 seeds within a white matter mask generated from 5-tissue-type segmentation of the T1-weighted images using MRtrix3 and FSL tools. The Frontal Aslant Tract (FAT) was automatically segmented and extracted using the FSL XTRACT atlas, employing seed, target1, and target2 regions of interest (ROIs) in the MRtrix3 tckgen command.

### 2.5. FAT dissection

After reconstructing the full FAT, we used TrackVis and Tractome toolkits to manually remove false-positive streamlines (Porro-Mun oz et al., 2015). The tract was then divided into three equal sub-bundles (posterior, middle and anterior) following the dissection approach outlined by Tagliaferri et al. (2023) and based on the anticipated spatial granularity of functional specialization in the superior frontal gyrus (SFG) (Cattaneo et al., 2018) and confirmed by anatomical characterization of the FAT in (Tagliaferri et al. 2024). The 3 dorsal (SFG) cortical origins of these sub-bundles served as targets for neuronavigated TMS, utilizing a neuronavigation system (SofTaxic software v3.8, EMS, Italy, connected to a Polaris infrared camera).

### 2.6. TMS session

TMS was administered using a MagPro 100 stimulator connected to a figure-of-eight coil (model MagPro MCF-B65) with each winding measuring 65 mm in diameter. The individual resting motor threshold (rMTh) was determined by stimulating the optimal spot over the left or right primary motor cortex (M1) that elicited a visible contraction in intrinsic hand muscles. The rMTh was defined as the minimum stimulation intensity that produced a visible hand muscle contraction in at least 50% of 10 consecutive trials. During the experiment, single-pulse TMS was delivered at an intensity set to 120% of each participant’s rMTh. TMS (120% of resting motor threshold) was delivered randomly in the 1.5-2.5 seconds interval within the 4-seconds trial. TMS was aimed at 3 spots on the SFG convexity, on the cortex associated with the origin of the FAT that were renamed SFG1, SFG2 and SFG3 illustrated in ***Figure 1*** [note that these corresponded to the P01, P03 and P05 positions of (Tagliaferri et al. 2023)] identified on the neuronavigated T1-weighted images, ensuring a local error of less than 1.7 mm on the neuronavigation system. To maintain the induced electrical field approximately perpendicular to the main sulci in the stimulated area, the coil handle was oriented laterally at a 90° angle to the midline when stimulating the SFG. Sham stimulation was performed by tilting the coil 90° away from the scalp surface and applying the stimulus to SFG2.)

**Figure 1:**
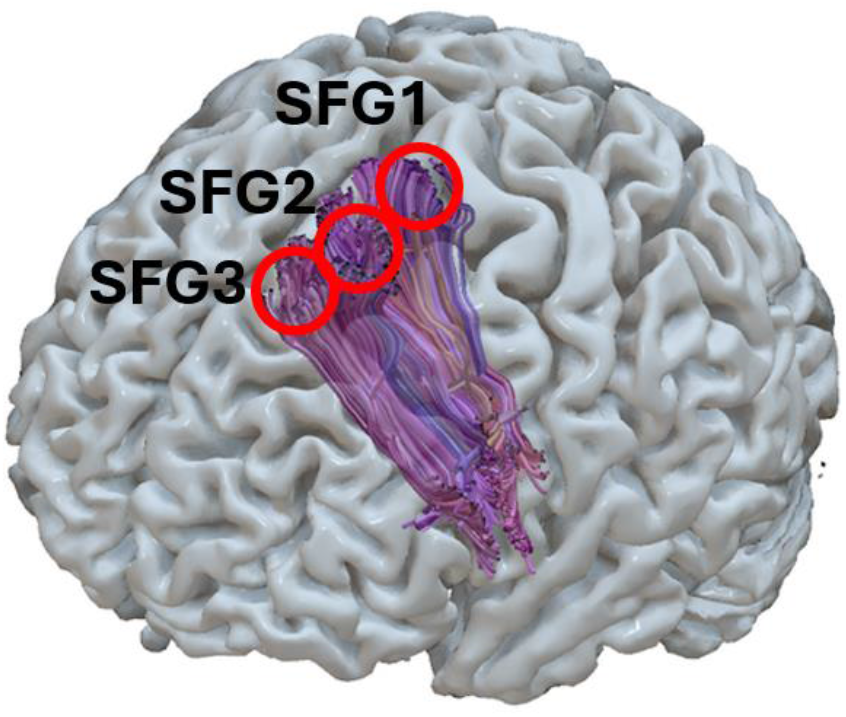
stimulation points along the origin of FAT fibers in the SFG-crown, divided into a posterior, middle and anterior bundle, in a representative subject

### 2.7. Experimental design

The experiment comprised two main tasks performed at each Frontal Aslant Tract (FAT) endpoint, with tasks alternated within each participant. The first task was a speech repetition exercise in which participants were instructed to repeatedly articulate the syllable “da” for 4 seconds, followed by a 2-second pause. This sequence was repeated to obtain a total of 30 audio recordings per stimulation point. Participants were encouraged to use their natural personal rhythm during the syllable repetitions. The second task involved a motor activity where participants tapped rhythmically on the spacebar of a keyboard continuously for 4 seconds. In each task, TMS was delivered in the middle of the 4 s sequence, with a +/-0.5 s jitter. For each FAT endpoint, the two tasks were alternated, and a sham condition was included as the third condition in a sequence of four conditions (comprising 3

FAT endpoints and one sham condition). Participants wore earplugs to minimize auditory distractions and had a microphone attached to their clothing to record audio data. The experimental procedures and data collection for both tasks were programmed and executed using E-Prime software.

### 2.8 Audio data preprocessing

The audio recordings collected during the speech repetition task were processed using a custom MATLAB script specifically designed for precise detection of speech onset times. This processing pipeline involved several critical stages: audio signal pre-processing, spectrogram analysis, onset detection, and interactive validation and correction through a graphical user interface (GUI). Initially, the raw audio signals were converted to mono by averaging the left and right channels, simplifying the data for analysis. To eliminate any potential DC offset that could bias the signal, the mean amplitude was subtracted from each recording. The signals were then normalized to have a maximum absolute amplitude of one, ensuring consistency across all recordings regardless of their original recording levels. To focus on the frequency components most relevant to human speech, the signals underwent band-pass filtering. A fourth-order Butterworth low-pass filter with a cutoff frequency of 1,000 Hz was applied to attenuate high-frequency noise above the typical speech frequency range. Additionally, a fourth-order high-pass filter with a cutoff frequency of 50 Hz was employed to reduce low-frequency disturbances such as background noise and recording artifacts. This filtering preserved the essential speech frequencies between 50 Hz and 1,000 Hz, encompassing fundamental frequencies and key formants. Spectrograms of the pre-processed signals were computed using the short-time Fourier transform (STFT) with a Hanning window of 256 samples and an overlap of 128 samples. This configuration provided a balance between temporal and spectral resolution, suitable for capturing the dynamics of speech. The mean power across selected frequency bins corresponding to the speech frequency range was calculated for each time frame, resulting in a time series representing the energy variations of the speech signal. The mean power time series was further smoothed using a Gaussian filter with a window size of 25 samples to reduce high-frequency fluctuations and enhance the detection of sustained increases in signal energy associated with speech onset. An adaptive threshold was established by smoothing the mean power with a larger Gaussian filter of 500 samples and scaling it by a threshold factor of 2. This adaptive approach allowed the threshold to adjust to local variations in the signal, improving the reliability of onset detection across different recordings and speaking styles. Onset points were identified by detecting moments where the smoothed mean power crossed above the adaptive threshold from below, indicating the initiation of speech. These preliminary detections were then subjected to user validation to ensure their accuracy. The timing of the syllables was then expressed in relation to the time of TMS (negative values before TMS and positive values after TMS.

### 2.9 Tapping data preprocessing

The data from the tapping task were collected using E-Prime software, which provided precise timing and recording of participants’ keystrokes during the task. Unlike the speech recordings, the tapping data did not require any signal preprocessing due to the discrete and digital nature of the keystroke events captured by the software. However, to maintain consistency in the analytical approach and facilitate direct comparisons between tasks, the tapping data were indexed and organized in a manner analogous to the speech onset times. The E-Prime software generated log files containing timestamps for each keystroke event, accurately reflecting the timing of participants’ taps throughout each trial. These raw timestamps were extracted and organized into a structured dataset using a custom script. For each trial, the keystroke times were indexed relative to the TMS pulse delivered at 2 seconds. The tap occurring nearest to the 2-second mark was assigned an index of zero. Taps preceding this reference tap were assigned negative indices, decrementing by one for each earlier tap (e.g., −1, −2, −3), while taps following the reference tap were assigned positive indices, incrementing by one for each subsequent tap (e.g., 1, 2, 3). This indexing mirrored the method used for the speech onset times.

### 2.10. Statistical analysis

Data analysis was based on a probabilistic account of the occurrence of motor events in the peri-TMS time interval. All the processes are outlined in Figure 2. First of all, we digitalized the data from both tasks marking as events the syllable onsets (Figure 8) and the finger taps. We then aligned all events to the TMS pulse and divided the timeline in bins 200 ms wide, counting occurrences of motor events within 200 ms-wide bins. The raw count of events within each bin was baseline corrected by subtracting the average value of the 5 bins pre-TMS (baseline period). We considered for analysis only 3 bins: the one before TMS, and the 2 after TMS. We analysed the overall data by means of a 3-way ANOVA with the following factors: TASK (2 levels: SPEECH and TAPPING), TMS (4 levels: SFG1, SFG2, SFG3 and SHAM) and BIN (BIN0, BIN1 and BIN2).

**Figure 2.**
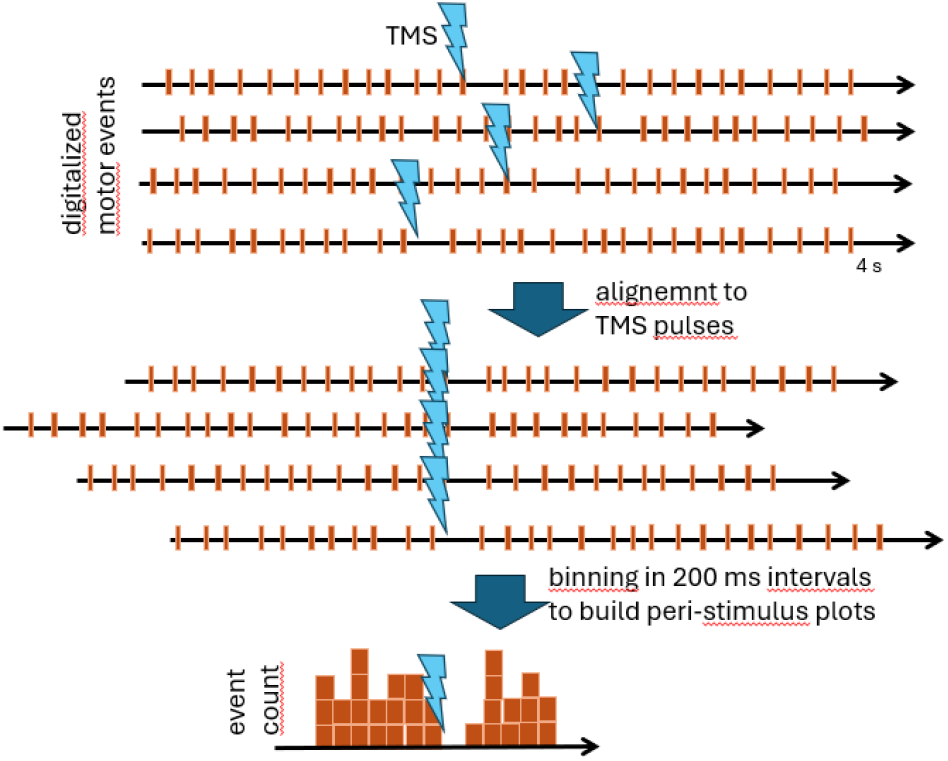
Steps of data pre-processing for peri-stimutus probability analysis

## 3. RESULTS

### 3.1 General results

None of the participants reported neither immediate nor delayed side effects of the procedure, which was generally well-tolerated as is usually the case for TMS of the vault of the scalp. A minority (3/21) of participants reported that TMS felt different in the sham condition.

### 3.2 Statistical results – omnibus TASK*TMNS*BIN ANOVA

We observed a significant TMS*BIN interaction [F(6, 120)=3.21, p=0.006, eta-squared=0.14, estimated alpha power=0.92] illustrated in ***Figure 3***.

**Figure 3:**
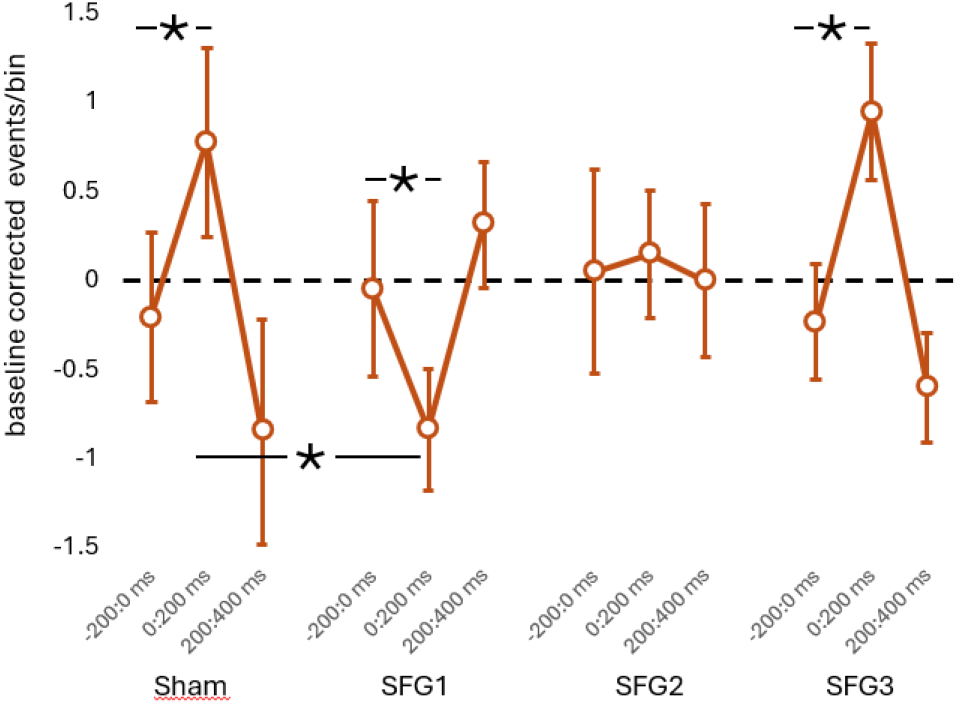
baseline-corrected, event count in the 3 bins(BIN0= −200:0 ms: BIN1-0:200 ms: BIN3= 200:400 ms) in teh Sham condition and in each of the 3 points of active stimulation. Asterisks indicate significant differences.

### 3.3. TASK*TMS sub-ANOVAs

Post-hoc analyses to explore the interaction were conducted by means of 3 separate 2-way ANOVAs (2*4 within-designs, with factors: TASK and TMS) on each of the 3 bins (in this way the possibility to test the comparison between active TMS and SHAM is preserved). Results showed no effect on BIN0, i.e. the pre TMS period: −200 to 0 ms (main effect of TMS: F(3, 60)=2.02, p=0.12), a significant main effect of TMS on BIN1, corresponding to the first 200 ms after TMS (F(3, 60)=4.68, p=0.005) and no main effect of TMS in BIN2, corresponding to the 200-400 ms period after TMS (F(3, 60)=1.78, p=0.16). Finally, the data from BIN1 were analysed by means of 3 t-tests for paired samples comparing Sham stimulation with the data from the 3 active spots and we found that only active stimulation on SFG01 was significantly different from sham (t(20)=2.78, p=0.011; all other p’s > 0.26 – note that threshold for significance is set to p=(0.05/3)=0.166 to account for the 3 multiple comparisons). The results therefore indicated that the main element driving the TMS*BIN interaction was specifically due to a difference between TMS over the SFG01 target compared to other conditions in BIN2.

### 3.4. TASK*BIN sub-ANOVAs

Finally, as a post-hoc measure, we explored the overall TMS*BIN interaction in a different way, by means of 4 separate two-way ANOVAs, one for each TMS condition, with 2 factors: TASK*BIN (2*3). The results showed a significant effect of BIN in the sham condition (F(2, 40)=3.37, p=0.044), a significant main effect of BIN for the SFG01 stimulation point (F(2, 40)= 3.99, p=0.026), no significant effect of BIN in the SFG2 (F(2, 40)=0.013, p=0.98) and a significant effect of BIN in the SFG3 spot (F(2, 40)=7.19, p=0.002). Post-hoc t-tests were conducted to evaluate the difference between BIN0 (baseline, pre-TMS) and the other 2 BINS. The results showed that in the Sham condition, TMS produced an increase in probability of motor events (t(20)=2.36, p=0.028) in the first 200 ms, compared to baseline (pre-TMS). In the SFG1 condition, TMS produced a significant decrease in probability of motor events in the first 200 ms after TMS, compared to pre-stimulation baseline (t(20)=-2.37, p=0.027). In the SFG2 spot, TMS we did not observe any differences between the bins and in the SFG3 we observed a pattern similar to that of Sham stimulation, i.e. TMS induced an increase in probability of firing in the 200 ms post stimulus (t(20)=3.04, p=0.006).

### 3.5. Summary of results

The full set of results is illustrated in ***Figure 3***. It is first worth noting that in none of the analyses the TASK factor contributed in any way to the variance of the data. Second, the TMS*BIN interaction indicated that TMS modulated the time-course of the peristimulus event plots differentially according to the stimulation condition. Interestingly we observed that Sham had an effect, i.e. that of increasing the probability of an event, shortly after stimulation. The same effect, indistinguishable form Sham was observed for stimulation over the SFG3, i.e. the most rostral spot. Stimulation over SFG1 (the most caudal spot produced an opposite pattern, with TMS inducing a decrease in probability of motor events in the first 200 ms. The intermediate point (SFG2) showed a pattern half-way between SFG1 and SFG3.

### 4. DISCUSSION

### Rhythmic motor patterns in the posterior FAT

We used tractography-guided TMS to target the posterior, middle and anterior origins of the fat in the SFG-crown cortex. We found that TMS to the posterior part of the SFG induced a drop in probability of producing a motor event, immediately after the TMS pulse. We have evidence that the neural representation is not that of actual movements, because it was independent from the effector used. Indeed, we found indistinguishable effects on similar actions performed with the distal upper limb (tapping) and with the orofacial system (syllable articulation). Given the spatial resolution of TMS (Cattaneo 2018), we cannot rule out between two possible contrasting hypotheses, that single neurons represent rhythm independently of the effector or whether within the stimulated region two populations of neurons individually represent either hand or mouth movements. In any of the two cases, we hypothesize therefore that the posterior SFG-crown contains the machinery for internal rhythmic movements, that is accessed and used whenever an internally-generated spontaneous motor pattern is desired. It is worth noting that we did not observe any significant rebound effect in the bin following BIN1 (i.e. the first post-TMS bin). This is finding is meaningful because the occurrence of a compensatory increase in frequency of motor events (rebound) following the observed decrease would have meant that rhythm had been reset in time by TMS. Thus, hypothesizing an interference with the actual clock function. On the contrary, the current data seem to point at the interference with the execution of a motor program that is clocked elsewhere (likely in the supplementary motor area (SMA))

### 4.1. Comparison with supplementary motor area (SMA) function

Interestingly, the capacity to perform rhythmic, spontaneous movements is commonly attributed to the SMA cortex which is known, mainly based on non-human primate data, to be able to calculate internally time intervals and to organize sequential movements, mainly by means of ramping neuronal activity (Macar et al. 2006; Casini and Vidal 2011; Cona and Semenza 2017). Our findings are not in contradiction with the main role of the SMA as internal clock for movement., On the contrary, we hypothesize that the crown of the SFG is part of a functional unit with the neighbouring SMA, receiving information from the SMA’s clock for goal-directed behaviour, and distributing it to different effectors. In humans, previous TMS studies have indicated that stimulation of the SMA produces alteration in spontaneous rhythmic behaviour similarly to the one observed here (Serrien et al. 2002; Steyvers et al. 2003; Schramm et al. 2019; Engelhardt et al. 2023; Kern et al. 2023). However, a careful look at the literature shows that in many of these studies, the actual sites of stimulation were so lateralized that the actual cortical site of stimulation was the SFG-crown and not the SMA. This is particularly evident in the works that used dense mapping of the cortical surface (Engelhardt et al. 2023; Kern et al. 2023), which clearly indicate that the local peak of effects on behaviour in the SFG-crown and not on the midline. In addition, some studies stimulating SMA on the midline used low stimulation intensities that are much more likely to affect the more superficial SFG-crown cortex rather than the deeper SMA (Serrien et al. 2002). Summing up, the comparison with the previous literature shows that the current results are fully compatible with the previous ones, and that it is likely that in many cases the main target of TMS was the SFG-crown besides the SMA-proper.

### 4.2 Hierarchical organization of the FAT

In our previous works exploring the FAT, we described how the middle portion of the FAT is involved in selection of strategies, when the choice is between anticipating or waiting for a given GO-signal (Tagliaferri et al. 2023). This conclusion was reached by observing the biases in strategy choices induced by TMS applied to the SFG and IFG origins of the FAT. Interestingly, owing to tractographic guiding of TMS, the cortical targets of the previous work are directly comparable to those of the present work. The SFG1 spot that was effective in the current experiment on interfering with action execution, is posterior to the SFG2 spot that showed an effect in the previous experiment on action choices. We show a hierarchical organization of the FAT and its associated cortex. We have previously demonstrated (Tagliaferri et al. 2023, 2024) that a) the FAT is anatomically organized in relatively independent, parallel modules arranged in a caudal-cranial order and b) that the mid-portion of the FAT is involved in strategy selection, by mediating competition between mutually-incompatible internally-timed and generated vs. externally-triggered behavior. Accordingly, we show here that the posterior portion of the FAT is involved in the direct production of internally-generated movements, confirming the hypothesis of a gradient of motor representations, form movements to actions and executive control. In addition, we confirm here, that individual information on anatomical connectivity (tractography) coupled with TMS can significantly increase the signal/noise ratio in spatial mapping of the cerebral cortex and provides a whole new way to interpret the functional mapping of the brain by non-invasive stimulation techniques.

